# Febuxostat enhances the anti-tumor efficacy of 2-fluoroadenine and 5’-methylthioadenosine in MTAP-deleted cancer

**DOI:** 10.64898/2026.05.19.726298

**Authors:** Baiqing Tang, Hyung-Ok Lee, Daniel Krzikike, Kathy Q. Cai, Sapna Gupta, Warren D. Kruger

## Abstract

**Background:** Homozygous deletion of the methylthioadenosine phosphorylase (*MTAP*) gene is a frequent genetic alteration in cancer. MTAP, which creates adenine from 5’-methylthioadenosine (MTA), is constitutively expressed in all tissues throughout the body. Previously, we described a novel strategy to specifically target *MTAP*-deleted cancer cells by combining the antipurine prodrug 2-fluoroadenine (2FA) with MTA. *In vit*ro, this combination efficiently killed *MTAP-* cancer cells, but *in vivo* the combination was much less effective in vivo. Here, we explored the role of xanthine oxidase (XO) in this process.

**Materials and Methods:** Various combinations of 2FA, MTA, and the xanthine oxidase inhibitor febuxostat (FX) were tested in various cancer cell lines grown *in vitro* and in mice. LC-MS/MS was used to examine the levels and ratio of intracellular 2-FA-containing nucleotides compared to adenine-containing nucleotides.

**Results and conclusions:** The treatment of cells with 2FA+MTA *in vitro* resulted in much higher 2FANP/ANP ratios than the same treatment *in vivo*. The addition of XO to culture media *in vitro* effectively abolished the killing by 2FA, and this effect was fully reversed by the addition of febuxostat (FX), a xanthine oxidase inhibitor. In *vivo*, the addition of FX to 2FA results in increased cell killing and toxicity and a 1000% increase in the amount of 2FA converted to 2-FA-monophosphate (2FAMP). Xenograft studies using *MTAP-* HT1080 and MiaPaCa-2 cell lines have shown that a 2FA/MTA/FX cocktail can cause tumor regression *in vivo*. These studies suggest that the combination of 2FA/MTA/FX should be explored as a treatment for *MTAP-* cancer.

## Introduction

Methylthioadenosine phosphorylase (MTAP) is a key enzyme in the purine and methionine salvage pathways that converts the polyamine byproduct 5′-methylthioadenosine (MTA) into adenine and methylthioribose-1-phosphate. It is a so-called “housekeeping” enzyme that is expressed constitutively in all tissues (1). The gene encoding human *MTAP* is located on chromosome 9p21, approximately 80 kb from the *CDKN2A/ARF* region (2, 3). Importantly, homozygous deletion of *MTAP* is one of the most frequent genetic alterations observed in cancer, with approximately 12% of all tumors exhibiting *MTAP* loss (4). Homozygous *MTAP* loss is especially frequent in some of the most lethal cancers, including those of the brain, pancreas, esophagus, and lung (5). The high frequency of *MTAP* loss in tumor cells has led to much interest in the development of novel therapeutic strategies that specifically target *MTAP* deletions in cells (6).

Previous work from our lab explored the use of a two-drug combination to specifically target *MTAP* deleted cells (7). In this approach, we combined the potent non-specific cell killing base analog, 2-fluoroadenine (2FA), with the MTAP substrate MTA, which acts as a protecting agent for *MTAP+* cells, but not for *MTAP-* tumor cells. We refer to this approach as “MTAP-mediated chemo-protection strategy” (Figure 1A). In *MTAP+* normal tissue, MTA is cleaved to adenine, which in turn is converted to adenine monophosphate (AMP) by the action of the enzyme adenine phosphoribosyltransferase (APRT). 2FA is an analog of adenine, a prodrug that must be converted into 2FAMP by APRT to become cytotoxic. Once 2FAMP is formed, it can be further converted into 2FADP and 2FATP (8). When *MTAP+* cells are incubated with five or more molar excesses of MTA to 2FA, the adenine produced from MTA competes with 2FA for APRT binding, resulting in reduced production of 2FAMP. In contrast, in *MTAP-* tumor cells, MTA is not converted to adenine, resulting in no competition for APRT and a greater percentage of 2FA being converted to 2FAMP. In cell culture, the 2FA/MTA combination killed *MTAP-* cells very efficiently after 48 h, but at the same concentration, showed much less toxicity against isogenic *MTAP+* cells (Figure 1B-C). However, in mice, we found that the effectiveness of 2FA in killing tumor cells was significantly reduced, resulting in only a modest difference in tumor growth kinetics between *MTAP+* and *MTAP-* xenografts in 2FA/MTA-treated mice (7).

**Figure 1.**
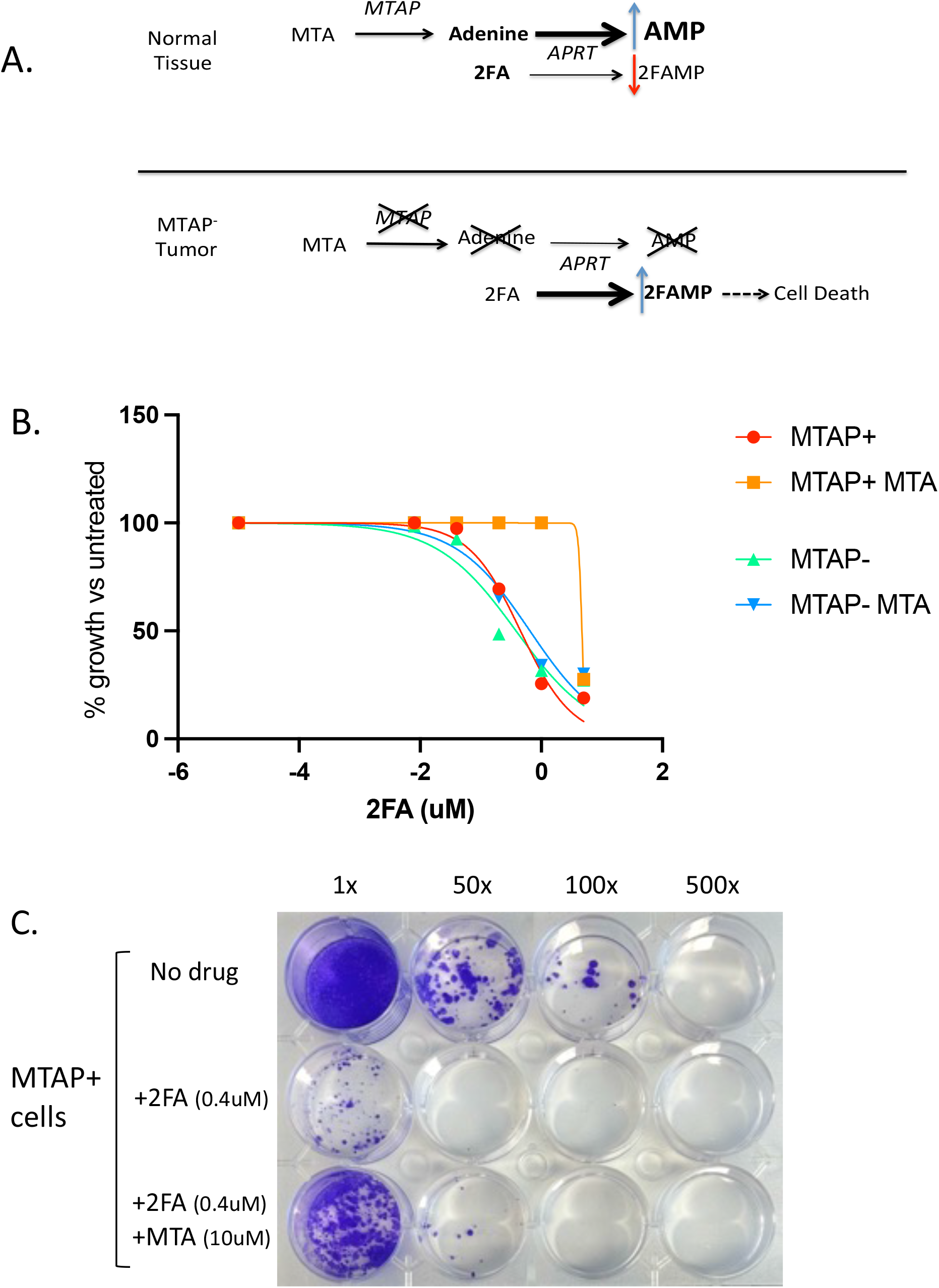
MTAP-mediated chemo-protection strategy. A. Scheme showing interaction between MTA and 2FA metabolism in *MTAP*+ and *MTAP*-cells. B. MTA protection in isogenic *MTAP*+ and *MTAP*-immortalized human pancreatic epithelial (HPNE) cells treated with 2FA in the presence or absence of 10 μM MTA for 48 hours. C. Colony formation showing MTA protection in mouse derived KPC3-3 pancreatic cancer (*MTAP*+) cells.

In the current study, we explored the hypothesis that the reduced effectiveness of 2FA in killing tumor cells *in vivo* may be due to the presence of the purine metabolic enzyme xanthine oxidase (XO). We showed that 2FA toxicity is modulated by XO and that the addition of an FDA-approved XO inhibitor greatly enhances the ability of 2FA/MTA to regress tumor growth *in vivo* in two different mouse xenograft models.

## Results

### 2FA incorporation in adenine nucleotides *in vitro* vs *in vivo*

To understand why 2FA/MTA was much more effective in killing *MTAP-* tumor cells *in vitro* than *in vivo*, we measured the ratio of 2FA-containing adenine nucleotide pools/adenine nucleotide pools in lysates from isogenic *MTAP+* and *MTAP-* HT1080 cells treated with either 2 μM 2FA alone or 2FA + 10 μM MTA (Figure 2A). In both *MTAP+* and *MTAP-* cells, 2FA treatment resulted in significant production of 2FA containing nucleotides (2FANP), with an overall concentration of 30-70% of the respective endogenous adenine nucleotide pools (ANP) at 24 h. When a 5-fold molar excess of MTA was added, we reduced the 2FANP/ANP ratio by more than 5-fold in MTAP+ cells, but did not affect MTAP-cells. These findings support the hypothesis that MTA inhibits the conversion of 2FA to 2FANP, and that an increase in the 2FANP/ANP ratio causes increased cell killing.

**Figure 2.**
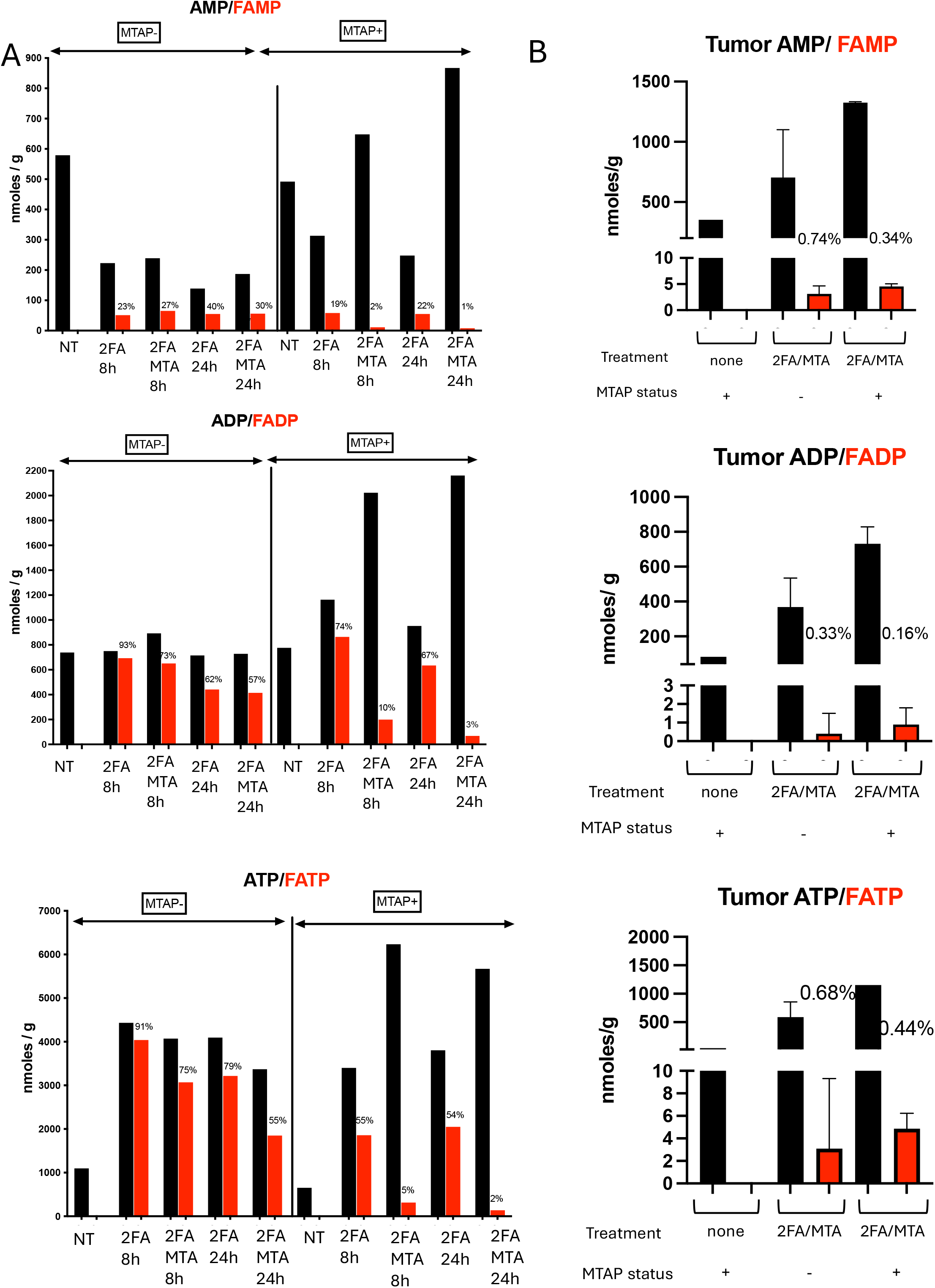
Nucleotide concentrations. A. Mono, di, and tri-nucleotide levels in isogenic MTAP+ and MTAP-HT1080 cells treated with either nothing (NT), 2 μM 2FA alone or with 10 μM MTA. Length of time of treatment was either 8 or 24 hours. Black bars show adenine nucleotides, while red bars show 2-flouroadenine (2FA) nucleotides. Percentage of 2FA relative to adenine for each nucleotide is shown above red bars. B. Nucleotide levels from HT1080 *MTAP+* and *MTAP-* tumors (n=3) grown SCID mice treated with either 2FA alone or 2FA+MTA. Error bars show SEM.

We next examined extracts derived from subcutaneous xenograft tumors of the same cell lines in mice treated with either nothing or four doses of 2FA/MTA (10 mg/kg 2FA, 50 mg/kg MTA) over an eight-day period (7). In tumors, we found that the concentration of 2FNP vs. ANP was less than 0.8% (Figure 2B) in both MTAP+ and MTAP-tumors. Importantly, the amount of 2FA administered to the mice was 10-times higher than that used in the cell culture experiment (making the highly conservative assumption that a mouse is 100% water). Taken together, these findings suggest that 2FA incorporation into nucleotide pools is far less efficient *in vivo* than *in vitro*.

### Xanthine Oxidase inhibits cell killing by 2FA

Based on the above findings, we hypothesized that there may be specific factors *in vivo* that could reduce 2FA bioavailability, thereby explaining the low concentrations of 2FANP and the limited effectiveness of 2FA/MTA to inhibit *in vivo* tumor growth. Xanthine oxidase (XO) is a purine catabolic enzyme that catalyzes the oxidation of hypoxanthine and xanthine to uric acid. Clinically, XO inhibitors, such as allopurinol or febuxostat, are used to treat gout (9). In humans and mice, XO is found at high levels in the intestine and liver, but it is also found at significant levels in extracellular fluids, such as serum (10). Because XO is known to have relatively low substrate specificity and can oxidize adenine to form 2,8-dihydroxyadenine (11), we tested whether XO addition could protect cells against 2FA-mediated cell death. Addition of XO to the media effectively protected HT1080 (*MTAP+*), MiaPaCa-2 (*MTAP-*), and AsPC-1 (*MTAP-*) from 2FA killing *in vitro*. This protection was entirely reversed by the addition of the XO inhibitor febuxostat (FX) (Figure 3, Supplemental Figure 1). These results demonstrated that XO can efficiently protect cells from 2FA.

**Figure 3.**
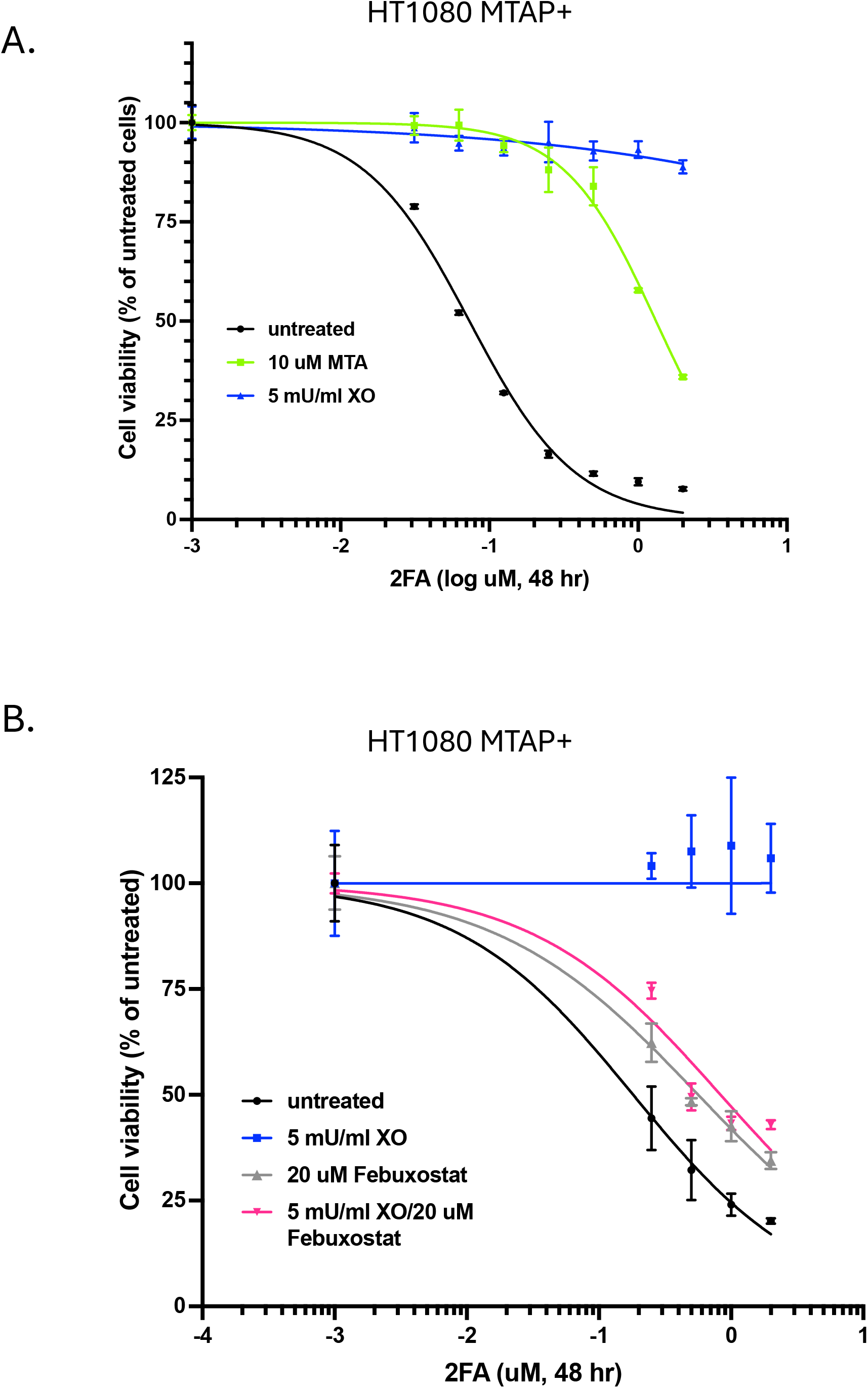
Xanthine oxidase inhibits 2FA toxicity. A. Growth inhibition of HT1080 *MTAP+* cells treated with various concentrations of 2FA for 48 hours in the presence or absence of either MTA or XO. B. Effect of XO inhibitor Febuxostat on XO protection. N=4 per point, error bars show SEM. Different letters next to points indicate significant difference.

### Effects of FX on 2FA toxicity in mice

To determine whether xanthine oxidase inhibition could enhance the in vivo activity of 2FA, we evaluated the combined toxicity of 2FA and febuxostat (FX) in mice. Animals were treated daily via oral gavage with 40 mg/kg 2FA, 5 mg/kg FX, or a combination of both. The body weight and clinical appearance were monitored throughout the experiment.

By day 4, mice receiving combination treatment exhibited marked lethargy and greater weight loss than those treated with 2FA or FX alone. Based on these observations, the dose of both agents was reduced by 25% in the combination group. Despite this adjustment, two mice in the 2FA/FX cohort were deceased on the morning of day 5 (Supplemental Video 1), prompting early termination of the study. Consistent with these clinical findings, animals treated with the drug combination showed significantly greater weight loss than the single-agent group (Figure 4A). Histopathological analysis further demonstrated increased tissue toxicity associated with the combination treatment (Figure 4B). Examination of the bone marrow, spleen, and colon revealed pronounced cellular depletion and architectural disruption in 2FA/FX-treated mice, whereas tissues from animals treated with 2FA or FX alone showed only mild or no abnormalities.

**Figure 4.**
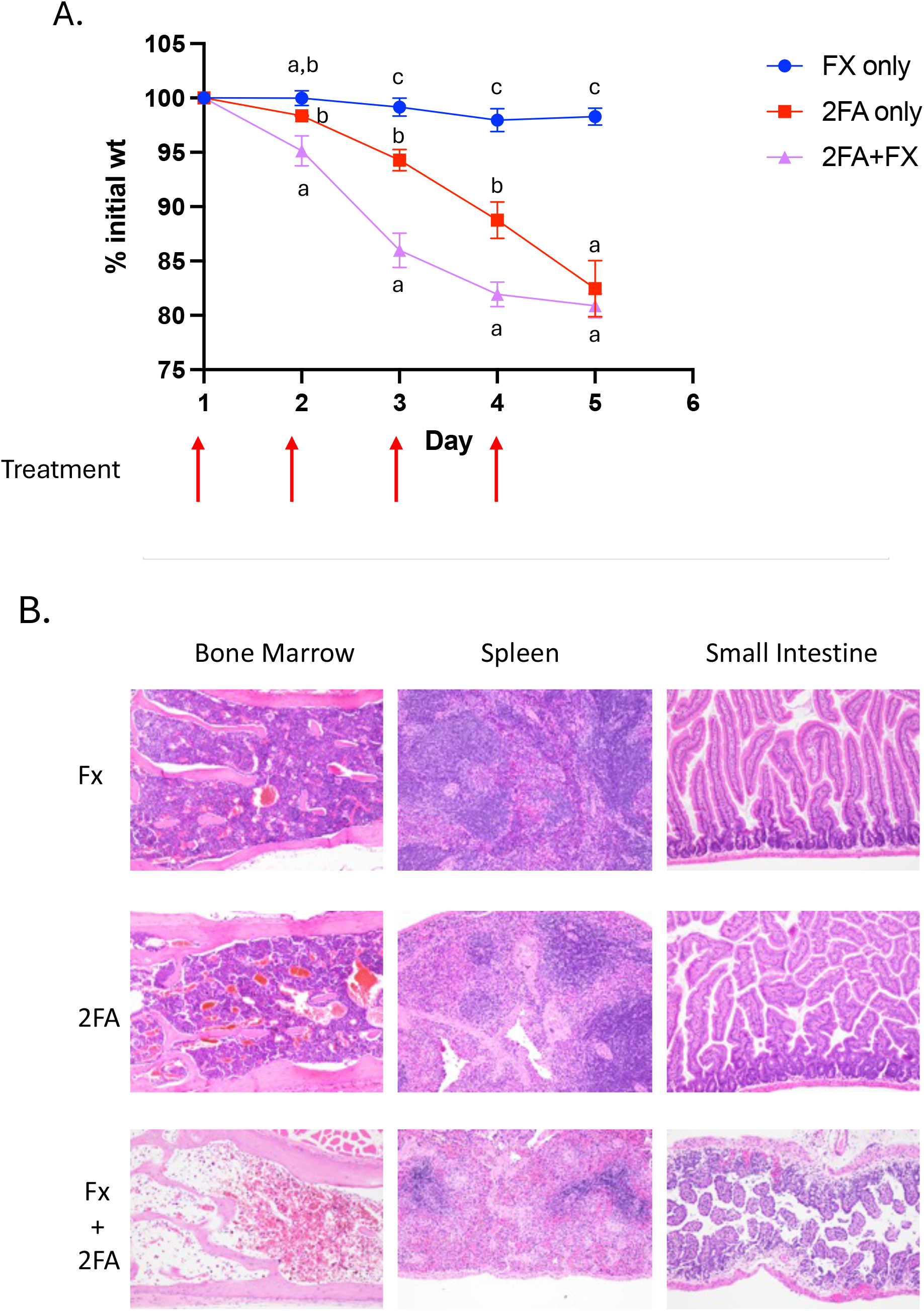
Febuxostat increases 2FA tissue toxicity in mice. Male 15-week-old C57BL6 mice were given orally either 40 mg/kg 2FA alone, 5 mg/kg FX alone, or both (N=4). Up arrows indicate treatment days. A. Percent body weight change in mice treated with indicated drugs. Different letters next to points indicate significant difference. B. Representative images of spleen, bone marrow, and small intestine of different groups (160x magnification).

We also examined the effect of FX on 2FAMP formation *in vivo* using a separate experiment. Mice were treated with vehicle, 2FA, 2FA/FX, or 2FA/MTA/FX for two consecutive days, and the pancreas and liver were harvested on the third day. The tissues were then assessed for 2FAMP and AMP concentrations (Figure 5). The addition of FX to 2FA resulted in a 1000% increase in the 2FAMP/AMP ratio in both the pancreas and the liver. Upon the addition of MTA, these ratios decreased by 75%. These studies showed that FX enhances and MTA suppresses the conversion of 2FA to 2FAMP.

**Figure 5.**
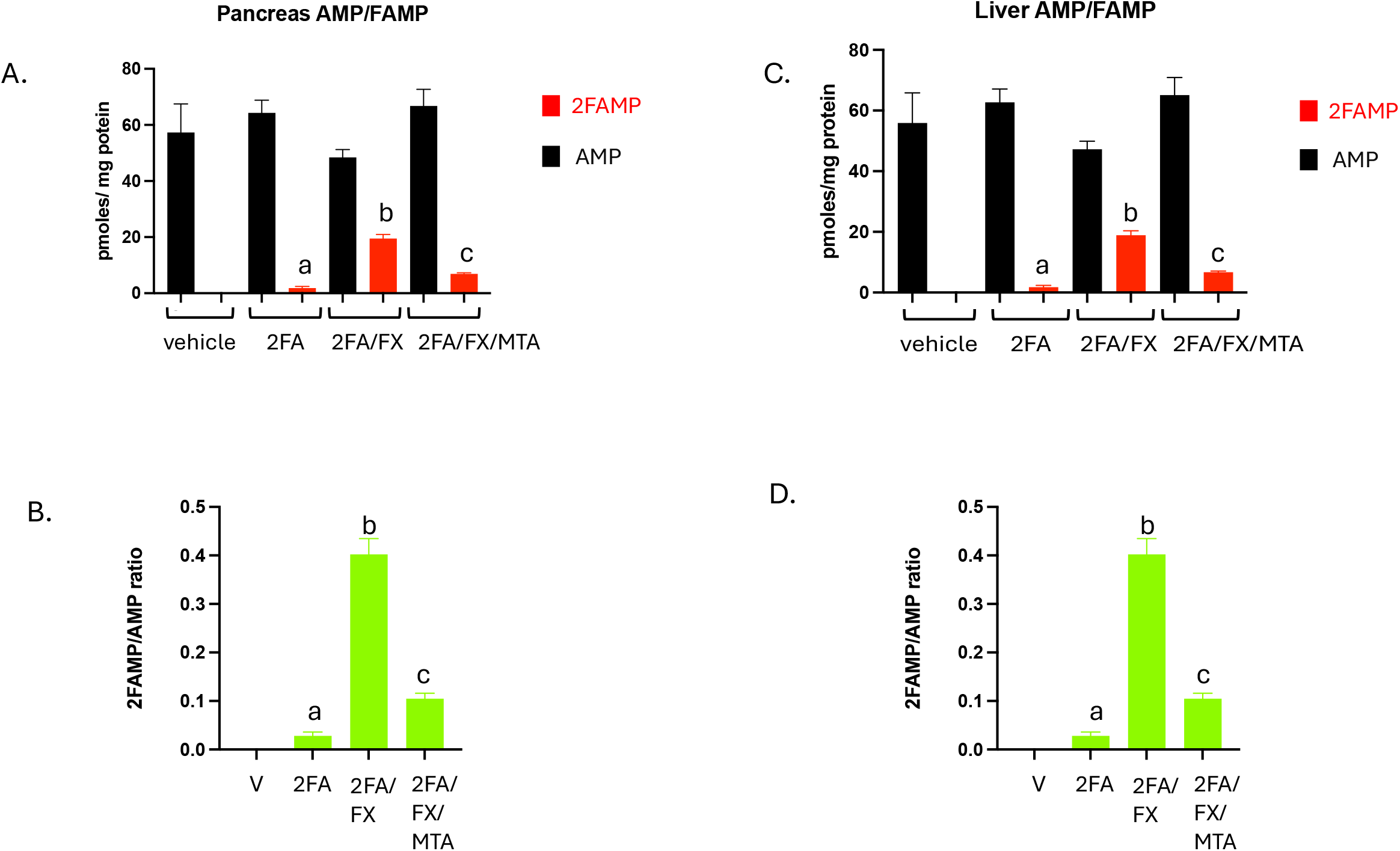
Tissue 2FAMP concentrations. A. Concentration of AMP and 2FAMP in pancreas tissue from mice given 2FA (30 mg/kg), 2FA+FX (3.75 mg/kg), 2FA+FX+MTA (180 mg/kg), or nothing. Mice were treated once per day for two days and tissue was harvested on day 3. N=3 per group, error bars show SEM. B. 2FAMP/AMP ratio for each group. C. Concentration of AMP and 2FAMP in liver. C. Ratio in liver. Different letters next to points indicate significant difference.

Together, these findings indicate that FX markedly enhances the systemic toxicity of 2FA *in vivo*. This increased toxicity is consistent with the hypothesis that xanthine oxidase normally limits 2FA bioavailability, and that its inhibition results in higher levels of active fluoroadenine metabolites in normal tissues.

### 2FA/FX/MTA inhibits xenograft tumor growth in immune compromised mice

We Next, we tested whether 2FA/MTA/FX improved tumor control in MTAP-deleted xenografts. First, we tested tumor growth inhibition using HT1080 *MTAP-* fibrosarcoma cells implanted in SCID mice. This model was identical to that used in our previous study (7), allowing us to determine whether the addition of FX increased the effectiveness of 2FA/MTA treatment. Tumor cells were injected into the flanks of 12 mice, and tumor size was determined after eight days. Mice were then divided into three groups (n=4) and treated with either vehicle (MTA/FX, 180 mg/kg/3.75 mg/kg), low dose 2FA/MTA/FX (10/90/3.75 mg/kg), or high dose 2FA/MTA/FX (20/180/3.75 mg/kg). A second dose was administered four days later, and on the eighth day, the mice were euthanized and tissue was collected. Tumors in the high-dose group exhibited a 60% reduction in size, whereas tumors in the vehicle group increased by 220% (Figure 6A). The tumors in the low-dose treatment group were slightly smaller than those in the vehicle-treated group, but this was not statistically significant. Tumors treated with high-dose 2FA/MTA/FX showed decreased levels of the proliferation marker KI-67 and increased levels of the apoptosis marker Caspase 3 and the inflammation marker myeloperoxidase (Figure 6B-C). These findings indicate that high-dose 2FA/MTA/FX can shrink HT1080 xenograft tumors and not simply slow growth. The toxicity of the treatment was assessed by monitoring the weight of the animals and histopathological examination of the thymus, spleen, intestine, and bone marrow at the end of the experiment. We observed increased weight loss and increased tissue toxicity in the 2FA-treated animals compared to those in the vehicle-treated animals (Supplemental Figure 2). This toxicity was more evident in the high-dose group.

**Figure 6.**
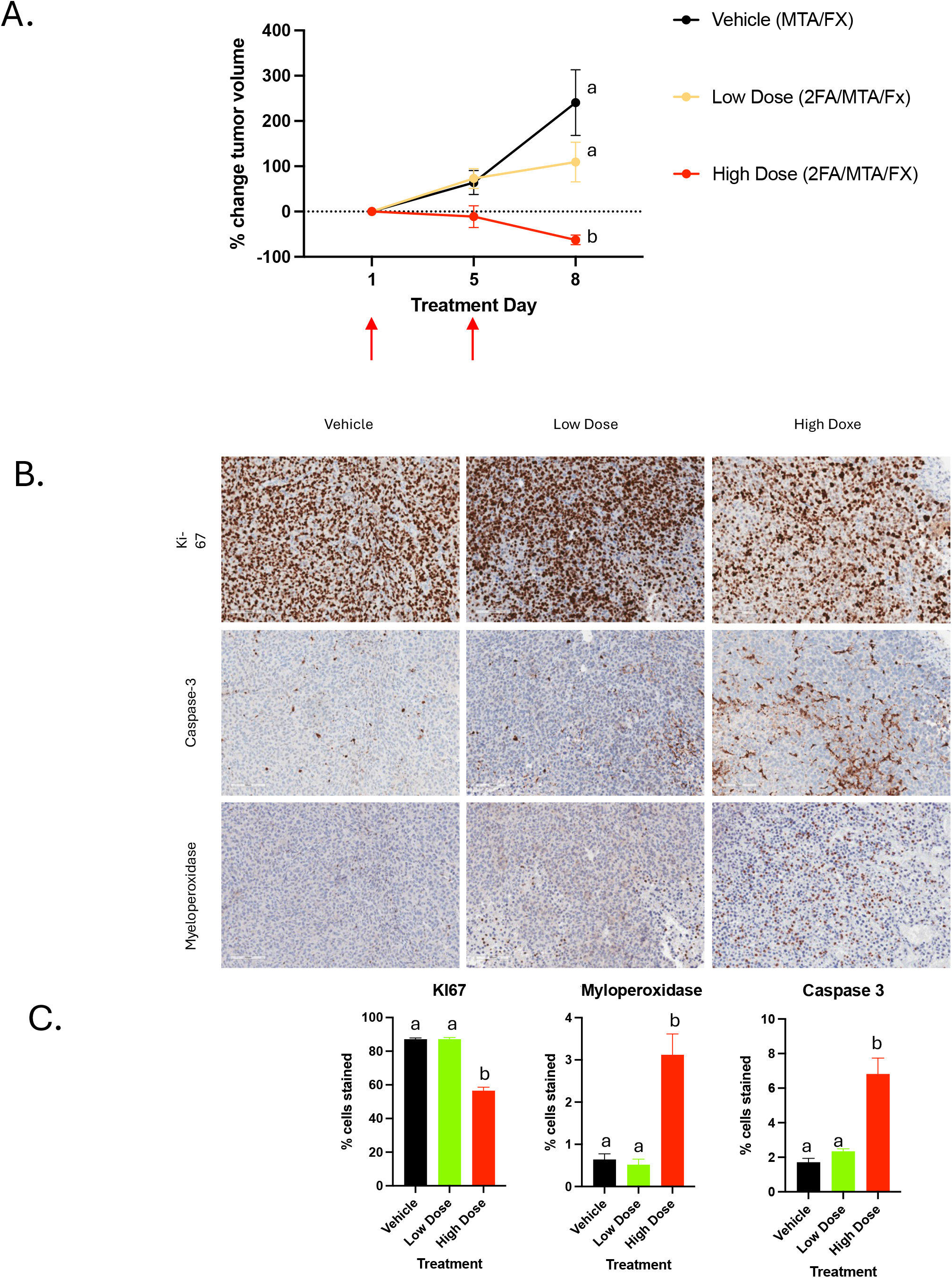
Growth of HT1080 cells in SCID mice. A. Average tumor volume in vehicle (no 2FA, 180 mg/kg MTA, 3.75 mg/kg FX), low dose treatment (10 mg/kg 2FA, 90 mg/kg MTA, 3.75 mg/kg FX) and high dose treatment (20 mg/kg 2FA, 180 mg/kg MTA). Arrows show days when drug was administered. N=4 per group. Note, one vehicle treated mouse was found dead on the day before the end of the study. B. Representative images of tumors stained for KI67, myeloperoxidase, and caspase 3. C. Quantitation of staining. Six to eight fields were quantitated for each sample.

The second and third experiments examined the effectiveness of 2FA/MTA/FX in treating tumors formed by MiaPaCa-2 cells, a *MTAP-* pancreatic adenocarcinoma derived line. The cells were implanted in 12 SCID mice and allowed to grow for one week. Unfortunately, during this week, one mouse had to be euthanized to fight the wounds. Mice were then treated with either vehicle (MTA/FX, 180 mg/kg/3.75 mg/kg) or two different doses of the 2FA/MTA/FX cocktail varying in the amount of 2FA but maintaining the 2FA/MTA ratio (10/180/3.75 mg/kg or 20/360/3.75 mg/kg). After the first dose, it appeared that the high-dose animals were suffering some toxicity, so all subsequent doses of 2FA concentration were halved. Tumors treated with high-dose 2FA/MTA/FX decreased in size during the treatment, and by the end of the treatment, was less than half the size of the tumors in untreated and low-dose mice (Figure 7A, Supplemental Figure 3A). During treatment, two animals (one in the high-dose group and one in the low-dose group) were judged morbid and euthanized before the end of the experiment. Both animals exhibited pronounced weight loss after the start of treatment (Supplemental Figure 3B). However, the other mice in both treatment groups showed either no or minimal weight loss; therefore, it was not clear whether these deaths were treatment-related, tumor-related, or due to other causes. H and E tissue analysis of the colon, liver, kidney, thymus, spleen, and bone marrow was performed on vehicle and high-dose mice at the end of the study. No significant differences were observed for any of the tissues, except for the bone marrow, where the one high-dose mouse that was euthanized early showed significant levels of toxicity (Supplemental Figure 3C).

**Figure 7.**
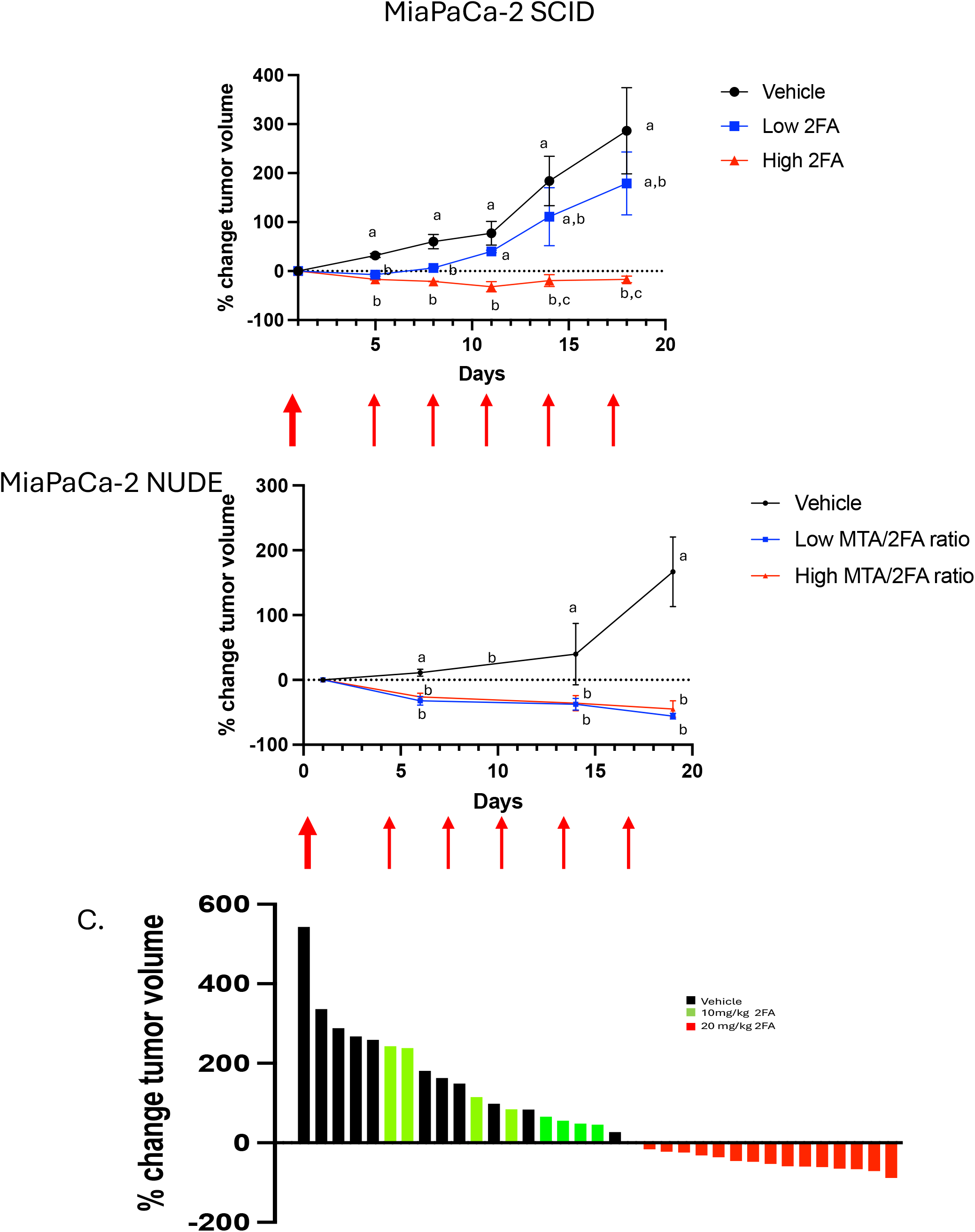
Growth of MiaPaCa-2 cells in mice. A. Tumor growth in SCID mice treated with either high 2FA mix (20 mg/kg 2FA, 180 mg/kg MTA, 3.75 mg/kg FX), low 2FA mix (10 mg/kg 2FA, 180 mg/kg MTA, 3.75 mg/kg FX) or vehicle (180 mg/kg MTA, 3.75 mg/kg FX). Arrows show time of treatment. After initial dose, concentrations of 2FA were halved for both the high and low dose groups. B. Tumor growth in Nude mice treated with either a low 2FA/MTA ratio (20 mg/kg 2FA, 180 mg/kg MTA, 3.75 mg/kg FX), or a high 2FA/MTA ratio (20 mg/kg 2FA, 360 mg/kg MTA, 3.75 mg/kg FX) or vehicle (180 mg/kg MTA, 3.75 mg/kg FX). Again, 2FA concentrations were halved after first dose. C. Waterfall plot for all xenografts combined.

Because three animals were lost during the experiment, we performed a second study using nude mice instead of SCID mice using the same cell line. The mice were treated with either vehicle (3.75 mg/kg FX, 180 mg/kg MTA) or two different ratios of 2FA/MTA (20/180 mg/kg or 20/360 mg/kg). As in the first experiment, the amount of 2FA was reduced by 50% after the first dose. Tumors in both treatment arms were significantly different from those in the vehicle arm, showing shrinkage compared to vehicle-treated animals (Figure 7b, Supplemental Figure 5A-B). However, reducing the 2FA/MTA ratio (i.e., increasing the amount of MTA) did not appear to offer benefits with regard to weight loss (Supplemental Figure 5C).

Taking the three different treatment experiments together, we found that all mice that were treated with at least one dose of 20 mg/kg 2FA FX showed tumor shrinkage during the experiment (Figure 7C). The mean tumor volume reduction in these animals was 47%, with 13/17 showing a 30% or greater reduction.

## Discussion

Homozygous deletion of the *MTAP* gene is one of the most frequent genetic rearrangements observed in cancer, with especially high rates in several tumor types and very low 5-year survival rates, including glioblastoma (loss rate 26-60%), pancreatic ductal adenocarcinoma (18-64%), cholangiocarcinoma (15%-30), esophageal cancer (12-27%), and non-small cell lung cancer (12-38%) (5). Therefore, targeting these alterations in cancer treatment is an area of intense interest. In our previous work, we developed a strategy in which we combined a “killing agent” (2FA) and an MTAP requiring a protecting agent (MTA) to specifically target *MTAP* deleted tumor cells (Figure 1A). In cell culture models, the 2FA/MTA combination was highly effective, but *in vivo* was less effective, only slowing MTAP-tumor growth and not actually reducing tumor size (7). Here, we showed that this difference in behavior was associated with a large difference in the amount of 2FA converted to 2FAMP in the two different systems. *In vitro*, the 2FAMP/AMP ratio is between 0.2 and 0.4, while *in vivo* the ratio was only about 0.0074, at least a 30-fold reduction. Given this finding, we hypothesized that 2FA is metabolized *in vivo* before being taken up by tumor cells. A likely candidate enzyme for this process is xanthine oxidase (XO), which can oxidize several purines, including xanthine, hypoxanthine, and adenine (12). XO is also known to be present at detectable levels in mouse serum (10). Consistent with this idea, we found that addition of the xanthine oxidase (XO) enzyme could reverse 2FA-induced cell death *in vitro* and that this effect could be reversed by the simultaneous addition of the specific XO inhibitor febuxostat (FX). To test the idea that XO in the serum or intestine might inactivate 2FA *in vivo*, we combined 2FA with FX and examined its toxicity in mice. We observed that FX addition substantially increased 2FA toxicity and resulted in the production of significantly more 2FAMP *in vivo*. In addition, we demonstrated that when MTA was added to the mixture, 2FAMP levels were reduced. Our findings showed that 2FA toxicity *in vivo* can be increased by FX and reduced by MTA.

Next, we performed a series of experiments to examine the effects of various concentrations of the 2FA/MTA/FX combination on three different *MTAP-* cancer xenograft models: mice treated with at least one dose of 20 mg/kg 2FA in combination with MTA and FX; 76% (13/17) mice showed at least a 30% decrease in tumor volume. However, we did not observe tumor regression using lower (10 mg/kg) doses of 2FA, suggesting that the treatment is dose-dependent. Overall, in comparison with our earlier studies using only 2FA/MTA, we found that the addition of FX significantly enhanced anti-tumor activity.

However, we found that 2FA/MTA/FX was more toxic. In all three treatment studies, the treated mice showed weight loss and some atrophy in the bone marrow and spleen compared to vehicle-treated mice. However, the level of toxicity varied among the mice. Although four mice were lost during the treatment stage, two of the four mice were treated with vehicle, suggesting that this was not due to 2FX toxicity. In our experience, immunocompromised animals implanted with tumors sometimes die unexpectedly, but we cannot rule out the possibility that these mice might have reacted to either MTA, FX, or perhaps the combination.

However, neither MTA nor FX have been shown to have significant toxicity in rodents near the concentration ranges used here (13-15). With regard to the optimal ratio of 2FA/MTA, we did not observe any difference in effectiveness between a 1/3.5 and a 1/7 molar ratio of 2FA/MTA, but the numbers were very small (n=4/group), which needs to be explored further. It is also important to point out that in this study, the cell lines we tested, HT1080 and MiaPaca-2, are both MTAP-; thus, we cannot be certain that the tumor regression we observed was specific to MTAP-cells. However, in our earlier studies (7), where we tested 2FA+MTA alone, we observed a strong effect of MTAP status on the growth inhibition of tumors.

Another approach to target MTAP loss involves the use of drugs that inhibit protein symmetric dimethyl arginine (sDMA) methylation by targeting PRMT5, the methyltransferase involved in this reaction. Second-generation PRMT5 inhibitors have been developed that cooperatively bind MTA to the PRMT5 active site and have shown promise in inhibiting the growth of *MTAP-* tumor cells in mouse models and in early human clinical trials. Recently, it has been demonstrated in a orthotopic mouse model of pancreatic cancer that combining a PRMT5 inhibitor with the DNA damaging agent gemcitabine (2’, 2’-difluoro 2’deoxycytidine) produced more tumor growth inhibition than gemcitabine alone (16). Given that gemcitabine and 2 fluoroadenine are both nucleoside analogs, it might be interesting to add a PRMT5 inhibitor to the 2FA/MTA/FX combination. As both treatments have increased specificity for *MTAP-* cells, the combination may have an even greater therapeutic window.

In summary, we showed that the addition of FX to 2FA/MTA enhances tumor cell killing *in vivo*. Given the high frequency of *MTAP* loss in various tumor types, we believe that this combination should be explored further.

## Methods

### Reagents

For cell culture studies, 2FA (Oakwood Chemical) and MTA (Sigma-Aldrich) were dissolved in DMSO and diluted in cell culture medium. 2FA was dissolved at a concentration of 0.36 mg/ml in the presence of equal normality sulfuric acid in a 1% carboxymethylcellulose solution. MTA was added to the 2FA solution at a concentration of 1.8 mg/ml. Xanthine Oxidase (Sigma-

Aldrich) was obtained as a lyophilized powder and was used at a concentration of 5 U/L (U= μmol xanthine oxidized/min). The febuxostat was obtained from obtained from Sigma-Aldrich.

### Cell line studies

Isogenic HT1080 MTAP+/− cells were created and maintained as previously described (17). KPC3-3 cells (*MTAP+*) were generated as previously described (18). MiaPaCa-2 cells were obtained from the ATCC. All cell lines were validated by STR analysis and tested for mycoplasma using a PCR Mycoplasma Detection Kit (Thermo Fisher). Isogenic MTAP+/-HPNE cells (19) were generated using CRISPR knockout, as previously described (7). The foci formation assay was performed as previously described (20). Drug sensitivity studies were performed using MTT assays 48 h after drug exposure, as previously described (7).

### LC MS/MS adenine and fluoroadenine nucleotide quantitation

Cell pellets were lysed in 500 µL of 12% perchloric acid, spiked with 10 nmol of ^15^N_5_-AMP (Sigma), and centrifuged to precipitate proteins. On the other hand, tumor tissues (Figure 2) were homogenized in 12% perchloric acid spiked with an internal standard to make a 10% homogenate, followed by centrifugation to precipitate protein. The supernatant was further neutralized with KOH to remove perchlorate by centrifugation, diluted to 2 mL with H2O, and then subjected to solid-phase extraction using a 1 cc Oasis WAX cartridge (Waters). The solid-phase extraction cartridges were conditioned with methanol and equilibrated with 50 mM ammonium acetate (pH 4.5). After the samples were loaded, the SPE cartridges were washed with ammonium acetate and eluted with 0.8 ml of methanol: water: ammonium hydroxide (80:15:5). The elution was then dried in SpeedVac and resuspended in a 50:50 water:acetonitrile solution. After drying and reconstitution, the samples were analyzed by LC-MS/MS using a Waters Acquity I-Class UPLC coupled to a Waters Xevo TQ-S Micro tandem quadrupole mass spectrometer. Chromatographic separation was achieved using a Waters Acquity BEH Amide column (2.1 × 150 mm, 1.7μm) at a flow rate of 0.2mL/min using a linear gradient from 80% to 70% mobile phase A over 12 min at 45°C. Mobile phase A consisted of 15 mM ammonium acetate and 0.3% ammonia in 90% acetonitrile, and mobile phase B consisted of 15 mM ammonium acetate and 0.3% ammonia in water (all solvents were Optima LC–MS grade, Fisher Scientific). For MS/MS detection, analytes were monitored using negative mode electrospray ionization with the following MRM transitions: AMP, m/z 346.03 → 78.87, ADP m/z 425.97 → 133.92, ATP m/z 505.97 → 158.83, 2FAMP 364.03 → 131.92, 2FADP 443.97 → 151.92, 2FATP 523.97 → 158.83, and N^15^-AMP 351.01 → 78.87. LC-MS/MS data were processed using the Waters MassLynx 4.2. Control experiments examining the linearity and sensitivity of the assay showed that the assay was linear (R^2^>.98) for all metabolites tested between 2 and 2000 pmol. The solid-phase extraction step was omitted from the mouse tissue experiments shown in Figure 5; therefore, only the monophosphates (2FAMP and AMP) could be quantified.

### Mouse studies

A summary of the six different mouse studies performed is shown in Supplemental data X. C57BL6 and SCID (C.B-*Igh-1*^*b*^/IcrTac-*Prkdc*^*scid*^) mice were obtained from Taconic Biosciences, while nude mice (NU/J) were obtained from Jackson Labs. All studies were reviewed and approved in compliance with the Fox Chase Cancer Center IACUC protocol #22-12. Mice were administered drugs by oral gavage at different concentrations and timings, as indicated in the figure legends. At the end of the experiments, the mice were euthanized, and tissues were collected, fixed in buffered formalin, and embedded in paraffin. Paraffin blocks were cut into 5-µm-thick sections that were placed on positively charged slides. Sections were stained with hematoxylin and eosin (HE) and mounted with Permount (Fischer Scientific, Pittsburgh, PA). For Caspase 3, Ki67, and myeloperoxidase, immunohistochemical staining was performed as described previously (7).

Either SCID or nude mice were injected into the hind flank with 2 × 10^7^ tumor cells in 200 μl of Matrigel. After seven days, the tumor size was measured with calipers (21), and mice were randomly divided into vehicle and treatment groups. Mice were then treated with either the vehicle or drug combination at the schedule indicated in the figures. Tumor growth was calculated by comparing the difference in volume at T_n_ with the volume at the start of treatment (T_0_).

Mice were euthanized by isoflurane overdose followed by cervical dislocation in accordance with the IACUC guidelines.

### Statistics

Pairwise comparisons between groups were performed using Student’s two-sided t-test. A p-value of 0.05 or less was considered significant. For tumor growth experiments, a single-sided ANOVA was used, followed by Dunnett’s post-hoc test against vehicle treatment at each time point. All graphs show the standard error of the mean (SEM). Different letters on the graphs indicate significant differences in the post hoc test. The number of replicates used for each experiment is shown in the appropriate figure legends.

## Data Availability

The data generated in this study are available upon request from the corresponding author.

## Acknowledgements

This work was funded in part by the National Institutes of NIH grant CA245871). We acknowledge the help of Fox Chase Cancer Center Laboratory Animal and Experimental Histopathology Facilities. We thank Dr. Paul Campbell for providing us with the HPNE cell line.

## Figure Legends

**Supplemental Figure 1**. Effect of XO and Febuxostat (FX) on 2FA toxicity at 48 h in two MTAP-deleted pancreatic carcinoma cell lines. N=4 per point.

**Supplemental Figure 2**. Weight loss and toxicity in HT1080 SCID mice. A. Graph of Weight Changes. B. Representative H and E staining of the thymus, spleen, small intestine, and bone marrow (160x).

**Supplemental Figure 3**. Analysis of the MiaPaCa-2 SCID experiment. A. Excised tumor weight. B. Individual mouse weight over time. C. Bone marrow H and E sections comparing vehicle- and high-dose 2FA-treated animals.

**Supplemental Figure 4**. Analysis of the MiaPaCa-2 nude mice A. Excised tumor weight. B Image of excised tumors. C. Changes in mouse weight during the experiment.

**Supplemental Video**. Animals on top were treated with FX only, those in the middle were treated with 2FA only, and those in the bottom were treated with 2FA+FX, as described in Figure 4.

