## Supplemental Figures for "Febuxostat enhances the anti-tumor efficacy of 2-fluoroadenine and 5’-methylthioadenosine in MTAP-deleted cancer"

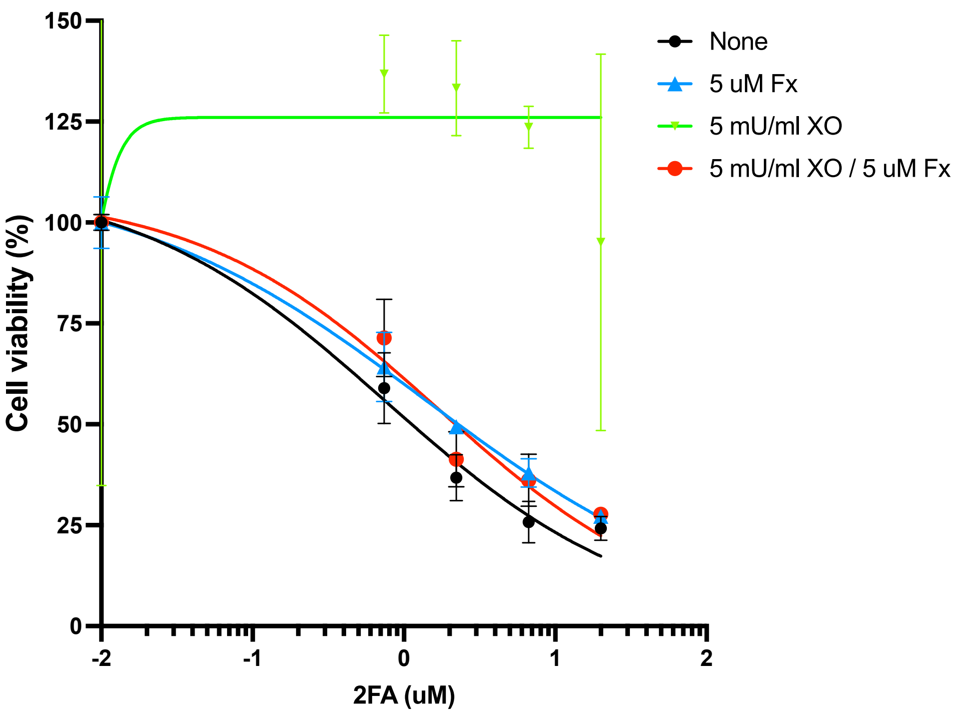


AsPC-1

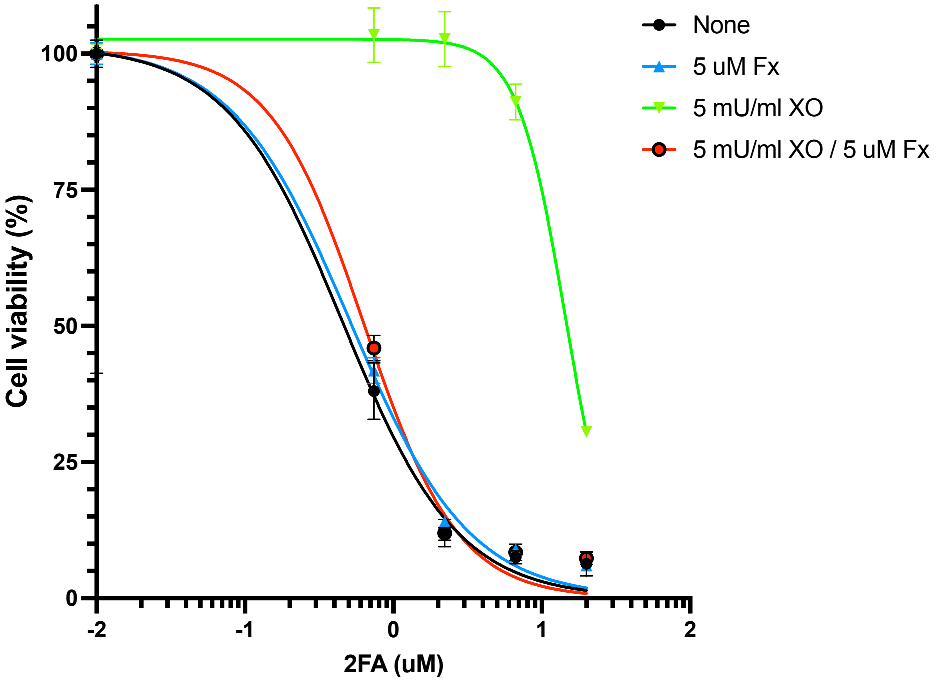


MiaPaca-2

Supplemental Figure 1

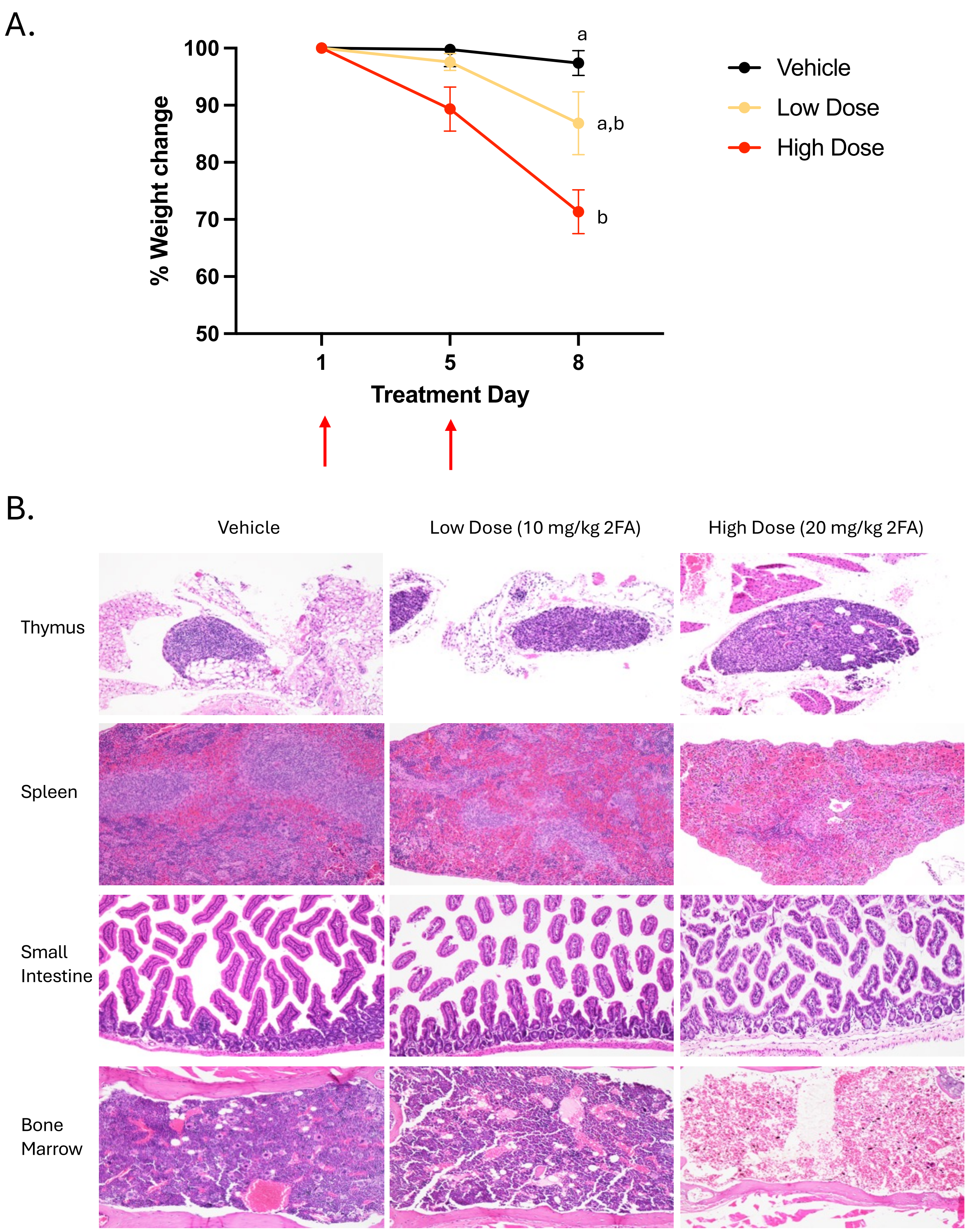


Supplemental Figure 2

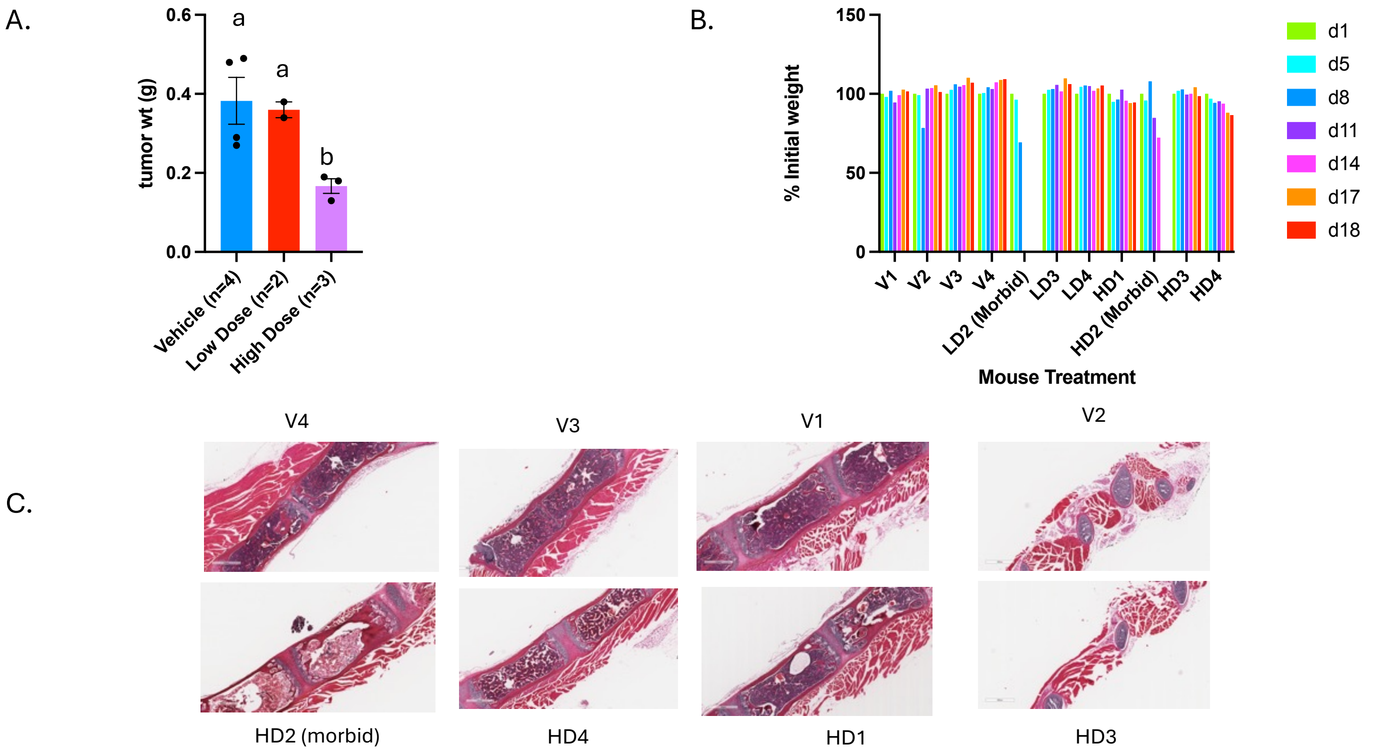


Supplemental Figure 3

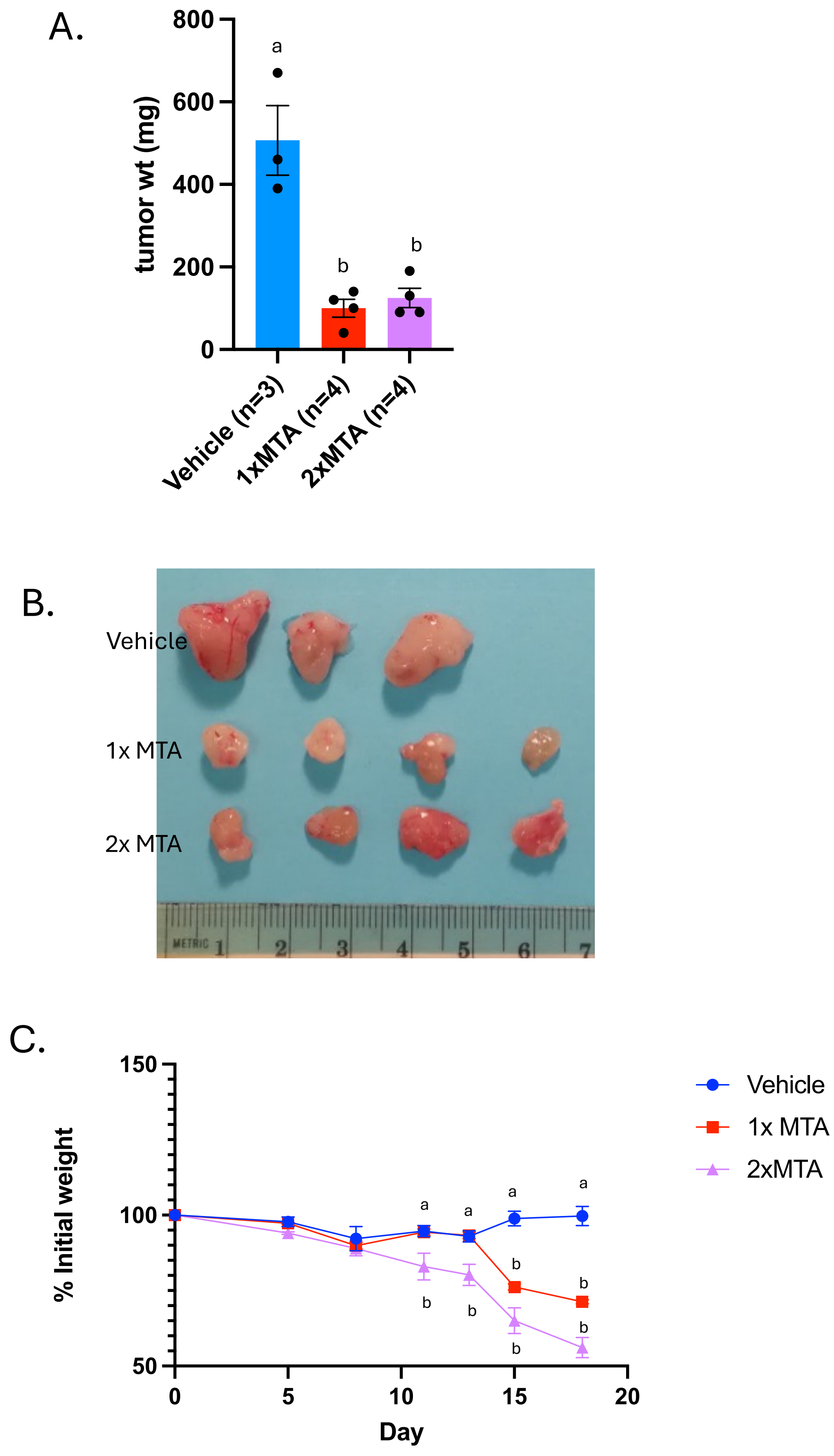


Supplemental Figure 4
